# ADP-dependent platelet activation is required for thrombus formation during a long-distance flight

**DOI:** 10.1101/2024.03.01.582882

**Authors:** Julie Tourn, Estelle Carminita, Lydie Crescence, Laurie Bruzzese, Nabil Adjriou, Regis Guieu, Christophe Dubois, Laurence Panicot-Dubois

## Abstract

The association between venous thromboembolism (VTE) and air travel is well documented. Prolonged exposure to reduced atmospheric pressure and low oxygen levels during flights triggers coagulation disorders, representing the primary risk factor for Deep Vein Thrombosis (DVT), surpassing immobility. In our study, we investigated how long-distance flight conditions affect VTE development in mice exposed to 6h of hypobaric hypoxia or normobaric normoxia after inferior vena cava (IVC) ligation. We observed a pro-thrombotic profile under flight-simulated conditions, characterized by larger thrombi with higher neutrophil and fibrin densities. However, no difference was observed in neutrophil extracellular traps (NETs) or fibrin-positive neutrophils in thrombi between groups, indicating that neutrophils/NETs may not be involved in DVT development during flight. Considering the elevated ADP levels observed at high altitudes, we further assessed thrombus formation in wild-type and *P_2_RY_12_*-deficient mice. Remarkably, thrombus formation was no longer affected by aircraft conditions in *P_2_RY_12_*-deficient or wild type mice treated with clopidogrel. We conclude that ADP-induced platelet activation is involved in the development of DVT during flight, suggesting that the use of P_2_RY_12_ inhibitors may be of interest to prevent DVT in susceptible patients.

## Introduction

Venous thromboembolism (VTE) is the third most common cardiovascular disease in the world after myocardial infarction and stroke, affecting nearly 10 million people per year (1). VTE includes DVT and pulmonary embolism (PE), the primary complication of DVT. Key factors in DVT development, known as the Virchow triad, include flow disturbance, endothelial activation or dysfunction and hypercoagulability (2). As venous stasis is the main trigger for this pathology, the thrombus primarily forms in vascular areas with a low shear rate, such as the venous valves of the lower limbs where the flow is reduced. Several risk factors for VTE have been identified, including cancer, genetic mutations, surgery, obesity, age (≥40 years), oral contraception, infections, prolonged immobility and long air travels (3). The association between air travel and VTE was first identified in 1977 by Symington and Stack as the “economy class syndrome” to describe the reduced space for leg extension typically found in economy class (4). Remaining seated for several hours in a restrictive space reduces venous blood flow by two-thirds, leading to the activation of the Virchow’s triad and thrombus formation (5). Based on these clinical reports, immobility has long been considered the main cause of VTE associated with long-distance flights. However, a crossover study conducted by Schreijer *et al.* revealed a hypercoagulable state after an 8-hour flight but no after an 8-hour movie marathon, indicating that the environmental conditions of the aircraft pose a greater risk of thrombosis than immobilization (5). Subsequent studies have shown that despite cabin pressurization, passengers are exposed to environmental conditions equivalent to an altitude between 1500-2400 meters (6). Prolonged exposure to reduced atmospheric pressure, and thus pO2, disrupts hemostasis and induces a shift toward a pro-coagulant profile. Several clinical studies have reported an increase in plasmatic levels of factor VIII, PAI-1, P-selectin, D-dimer or pro-inflammatory markers after a long-distance flight, indicating a defect in hemostasis (5,7). Interestingly, the activity of plasma factor VIIa is similar after a long-distance flight as following a movie marathon, indicating that the activation of the zymogen FVII is the results of a prolonged immobilization rather than aircraft conditions (8). Although monitoring coagulation and inflammation markers after exposure to aircraft conditions provides some insight into the development of a pro-coagulant profile, the mechanisms associated with thrombus formation during air travel remain poorly understood. With billions of air travelers each year, it is essential to identify the factors involved in coagulation disorders during flight.

In this study we recreated the environmental conditions characteristics of a long-distance air travel to examine the impacts of air-travel on DVT development in mice. We monitored the differences in size and composition at local level (thrombus) and at systemic level (blood compartment) due to hypobaric and hypoxia exposure. The participation of the coagulation cascade, NETs and the activation of platelets in the thrombus were evaluated. Finally, we focused on the P_2_RY_12_ pathway, which appears to be an important trigger for DVT occurring during long air travels. Collectively, our results indicate that targeting platelets, rather than the blood coagulation cascade, may constitute an alternative to prevent DVT during a long-distance flight.

## Results

### Aircraft conditions promote thrombus formation during DVT by acting both in circulation and locally

We first evaluated the effect of long-distance flight conditions on thrombus formation after 24 hours of IVC ligation. We observed that exposure to hypobaric hypoxia conditions increased the frequency of thrombus formation compared with standard environment, with 100% of mice that developed a thrombus versus 70% respectively **(Figure 1A)**. We conclude that mice exposed to flight conditions are more sensitive to stenosis triggered by IVC ligation. Morphological differences were observed between thrombi recovered under standard or hypobaric hypoxia conditions, with thrombi appearing larger after exposition to long-distance flight conditions (Figure 1B). The weight of thrombi formed in mice exposed to hypobaric hypoxia conditions was also higher compared to control group *(17,43±3,48 vs 6,322±2,868 mg)* **(Figures 1C)**. Furthermore, the surface area of thrombi harvested in the flight group was 12 times greater than in mice under standard conditions **(Figure 1D)**. These results indicate an induction of a pro-thrombotic profile in mice exposed to flight conditions that promote the incidence of venous thrombosis. Thrombus formation during long-distance flight simulation is enhanced or even accelerated, leading to the formation of denser thrombi. Aircraft conditions promote, therefore, thrombus formation in the flow restriction model of the IVC.

**Figure. 1.**
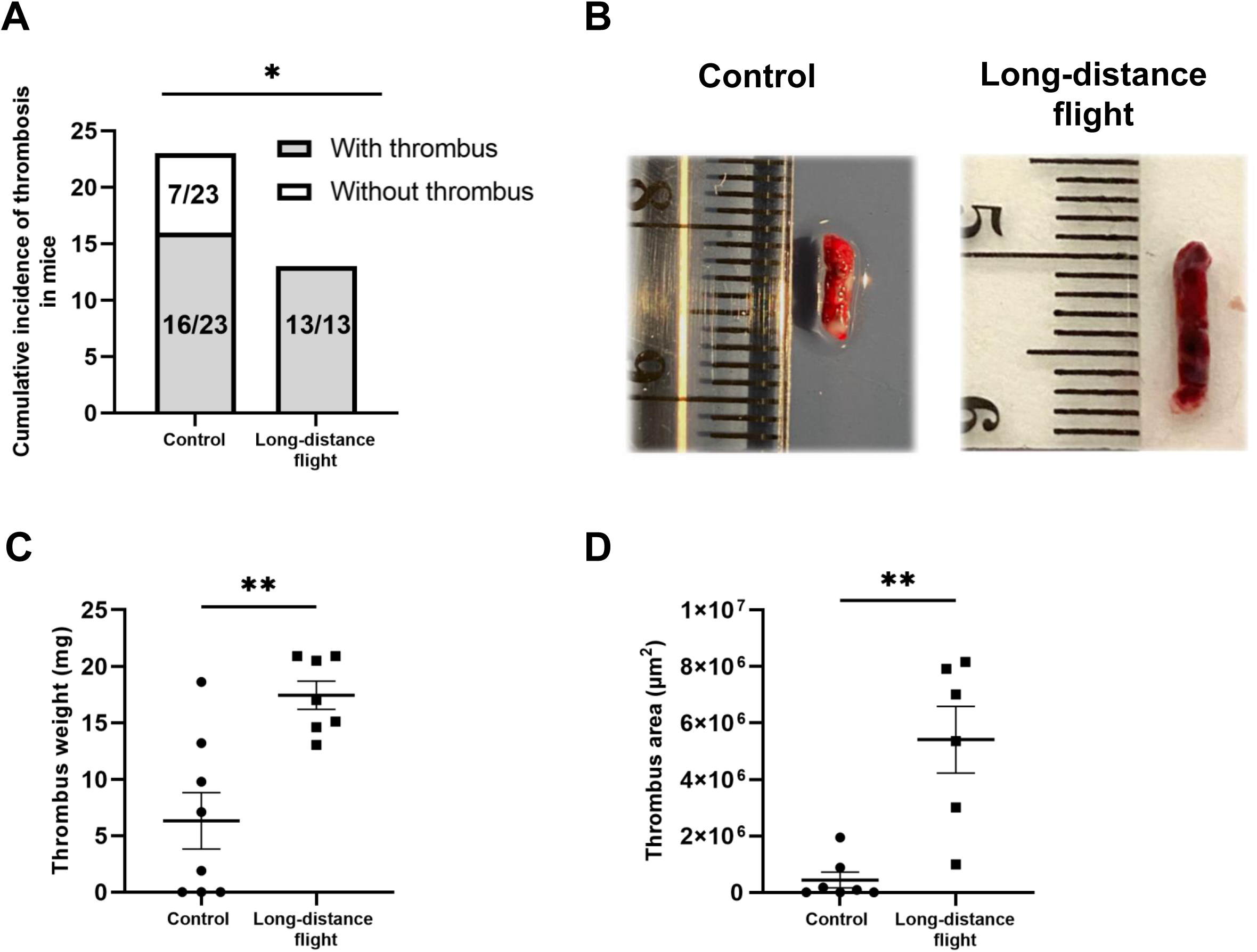
Long-distance flight conditions increase weight, area and frequency of thrombus formation after 24 hours of DVT. **(A)** The graph illustrates the aggregated occurrence of thrombosis in mice 24 hours following the induction of inferior vena cava (IVC) stenosis under normobaric normoxia (control) and hypobaric hypoxia (long-distance flight) conditions. (Mann-Whitney test, * p≤0,05). **(B)** Representative pictures of thrombi collected in wild type mice exposed to control or long-distance flight conditions. **(C-D)** Graphs represent weight **(C)** and area **(D)** of thrombi in mice exposed to control or long-distance flight conditions. (Student Test, ** p≤0,005).

To better understand the impact of long-distance flight on the development of DVT, we next investigated the effect of hypobaric hypoxia on the concentration of circulating cells after flow restriction in the IVC. Although no difference was observed in the concentration of circulating platelets, monocytes, and red blood cells, a significant decrease in the count of neutrophils and lymphocytes in the flight conditions group was determined **(Figures 2A-E)**. However, the expression of the activation markers, i.e. neutrophil elastase and the activated form of CD11b, was identical between the two groups of mice **(Figures 2F, 2G)**, indicating that circulating neutrophils were not activated under hypobaric hypoxia conditions. Consistent with these results, the prothrombin time was not impaired in mice exposed to the environmental conditions of the aircraft in comparison with the control group **(Figure 2H)**. These results suggest that part of circulating neutrophils and lymphocytes have mostly been consumed in the thrombus. Of note, the number of circulating leucocytes were also decreased as a result of IVC ligation in both environmental conditions, but in a greater extent in the flight group, suggesting that the reduction of the number of neutrophils and lymphocytes in mice with DVT was the result of both the surgery and hypobaric hypoxia (**Supl. Figure 1**). Nevertheless, the consumption of lymphocytes and neutrophils induced by inflammation following DVT is enhanced by aircraft conditions.

**Figure 2.**
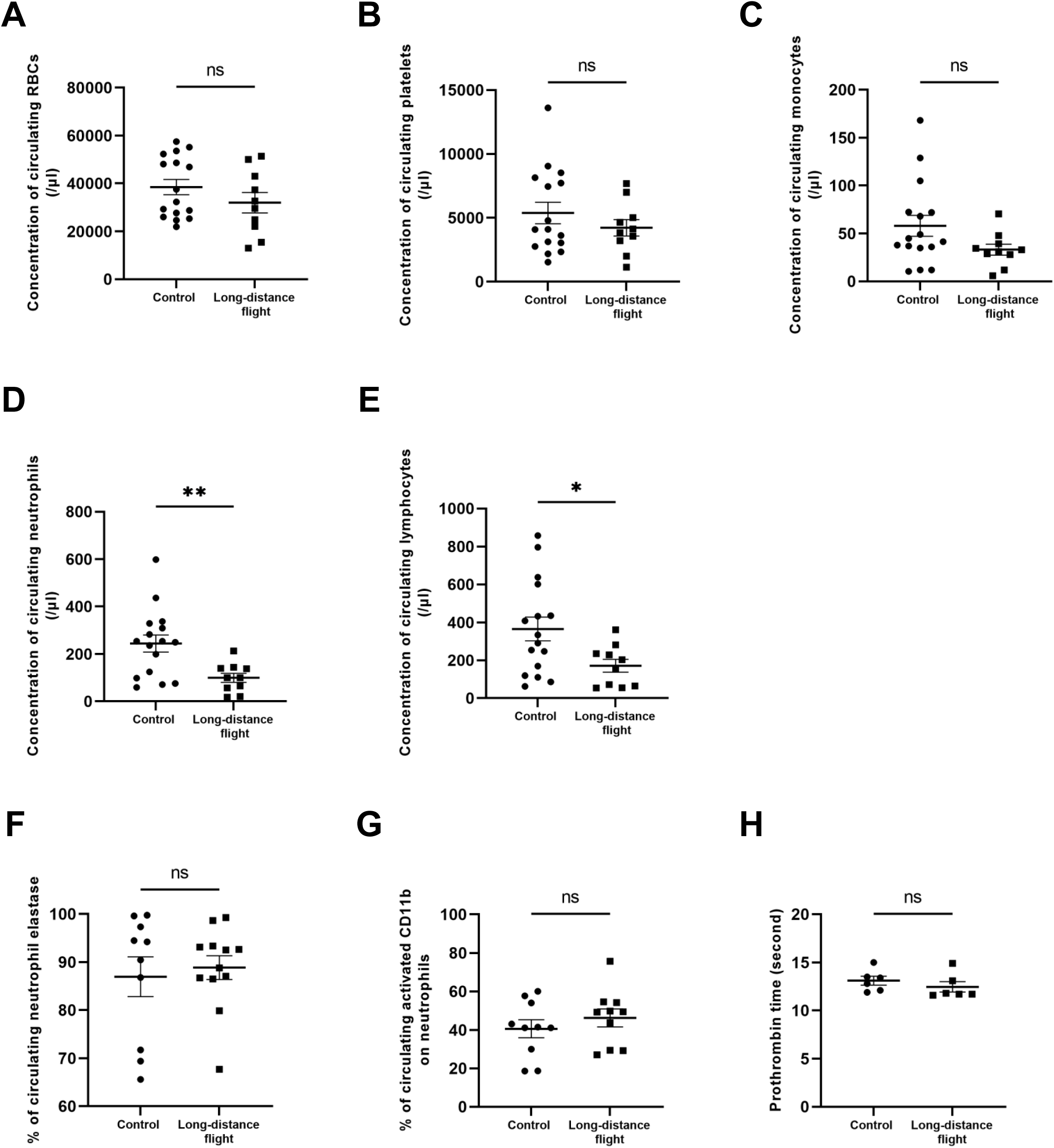
Long-distance flight conditions decrease the concentration of circulating leukocytes without leading to a pro-coagulant state. **(A-E)** Graphs depict the concentration of circulating erythrocytes **(A)**, platelets **(B)**, monocytes **(C)**, neutrophils **(D)** and lymphocytes **(E)** in mouse exposed to normobaric normoxia (control) and hypobaric hypoxia (long-distance flight) conditions. Twenty-four hours post-IVC stenosis, blood samples were collected, and the circulating concentration of cells was assessed using flow cytometry. Erythrocytes were identified using the anti-Ter119 antibody (0,5 µg/ml), platelets were labeled with the anti-X488 antibody (0,1 µg/ml), neutrophils with anti-Ly6G antibody (0,2 µg/ml), monocytes with anti-Ly6C antibody (0,2 µg/ml) and lymphocytes were detected according to the CD45^+^, Ly6G^-^ and Ly6C^-^ population. **(F-G)** Graph represents the percentage of neutrophil elastase **(F)** and the activated form of CD11b (0,2 µg/ml) **(G)** on circulating neutrophils in control or exposed mice. **(H)** Blood was collected from the tail to evaluate the pro-thrombin time in each group. (Student Test, * P≤0,05, **P≤0,005).

To define whether a long-distance flight altered the composition of the thrombus, we next compared, by immunofluorescence analysis, the composition of the different thrombi obtained. Whereas no difference was detected on platelets, a significant decreased was observed in the fluorescent signal corresponding to the presence of red blood cells by up to 36% after exposure to hypobaric hypoxia conditions compared to the control group **(Figures 3A and 3B)**. Also, the fluorescence intensities corresponding to the presence of leukocytes and neutrophils, were more than twice as high in thrombi coming from the flight group compared to mice remaining in standard conditions **(Figures 3C and 3D**), confirming that circulating neutrophils may be consumed in larger thrombi observed in the flight condition group.

**Figure 3.**
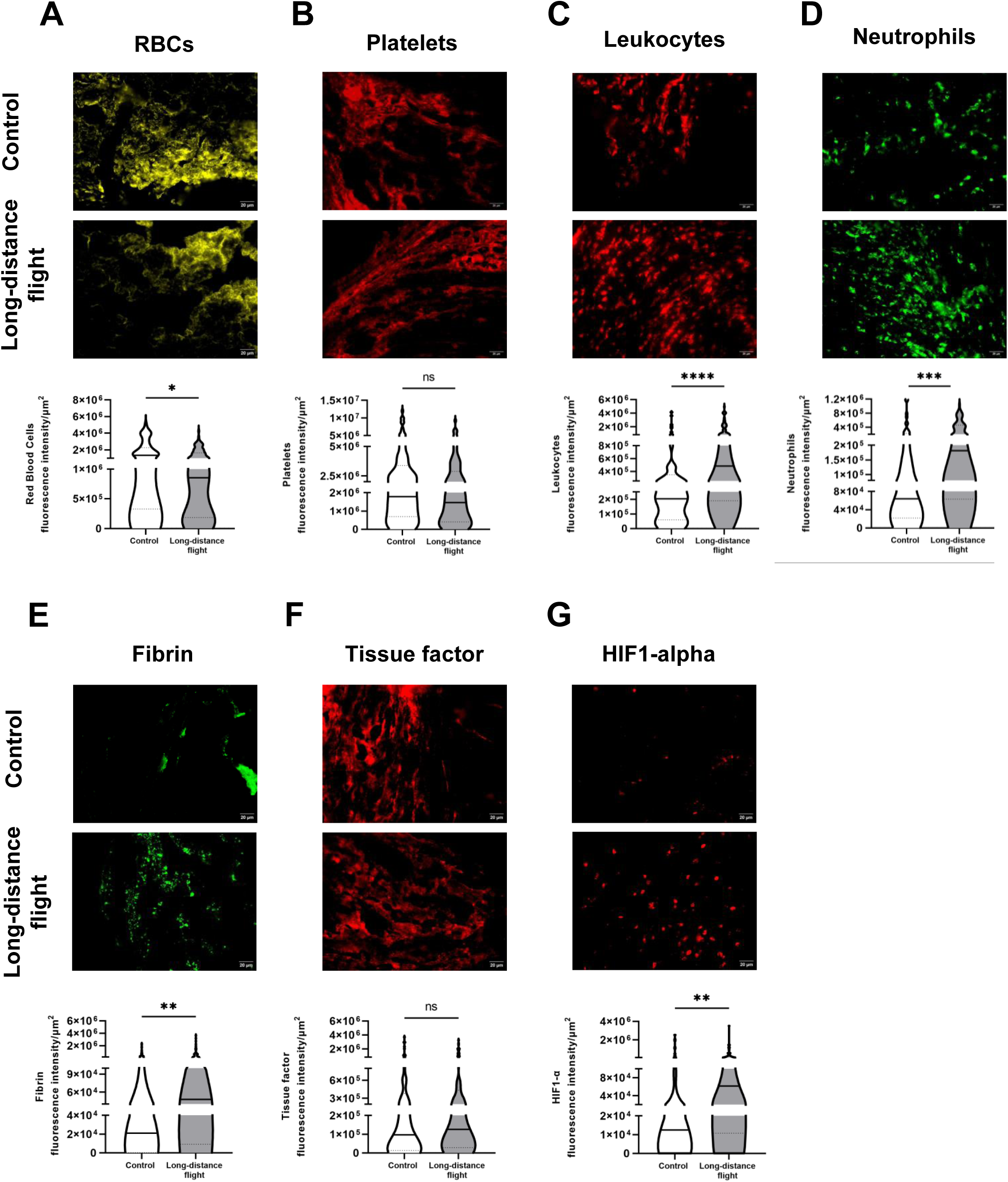
Long-distance flight conditions affect both the cellular and protein composition of thrombi. **(A-G)** Representative images captured with x40 objective, associated with graph, depict the fluorescence intensity of erythrocytes **(A)**, platelets **(B)**, leukocytes **(C)**, neutrophils **(D)**, fibrin **(E)**, tissue factor **(F)** and HIF1-α **(G)** in mouse exposed to normobaric normoxia (control) and hypobaric hypoxia (long-distance flight) conditions (n=5). Twenty-four hours post-IVC stenosis, thrombi were collected in both groups and fixed in OCT to perform cryostat sections (5 µm). Erythrocytes were immunoassayed with anti-Ter 119 antibody, platelets with anti-X647 antibody, leukocytes with anti-CD45 antibody, neutrophils with anti-Ly6G antibody, fibrin with anti-fibrin antibody, tissue factor with anti-tissue factor antibody and HIF1-alpha with anti-HIF1 alpha antibody. All antibodies were used at a final concentration of 1 µg/ml. Lines show the median. (Mann-Whitney test, * P≤0,05, ** P≤0,005, *** ≤0,0005, P**** P≤0,0005).

We next investigated the activation of the coagulation cascade through the generation of fibrin. The fluorescent signal corresponding to fibrin deposition was twice as important after exposure to aircraft conditions than the one observed in control mice **(Figure 3E)**, suggesting an over-activation of coagulation induced by hypobaric hypoxia conditions. However, in accordance with the literature (8), no difference in tissue factor expression was observed between the two groups **(Figure 3F)**. Finally, the fluorescent signal corresponding to the hypoxia inducible factor 1 (HIF1-alpha), which triggers the expression of pro-inflammatory and pro-thrombotic genes, was almost twice as high in thrombi of mice exposed to a long-distance flight **(Figure 3G)**. Long-distance flight conditions act, therefore, locally by increasing the deposition of pro-coagulants elements in the thrombus, independently of the extrinsic pathway of coagulation.

### Neutrophils and NETs are not involved in thrombus formation during long-distance flight

We next wondered whether neutrophil activation markers and NETs could be overexpressed during hypobaric hypoxia. By assessing the fluorescence intensity of citrullinated histone 3, an element of NETs structure, and Hoechst to identify the nuclei of cells, we did not observe strings of DNA in the thrombus or any difference in the expression of CitH3 between both groups (control and long-distance flight) **(Figures 4A and 4B)**. In accordance with our in vivo descriptions, we did not detect the presence of NETs on purified murine neutrophils chemically treated with increasing concentrations of cobalt chloride (CoCl_2_) **(Figures 5A-D)** or cultured in a hypoxia chamber **(Figures 5E and 5F)**. These results indicate that hypobaric hypoxia doesn’t stimulate NETs production, suggesting that the pro-thrombotic phenotype observed during flight is independent of NETosis. In addition, neutrophils in thrombi developed in mice exposed to aircraft conditions do not tend to be more activated based on the CitH3 signal. Although they do not form NETs, confocal microscopy revealed that neutrophils colocalized with fibrin **(Supl. Figures 2A-C)**. Of note, the colocalization of the fluorescent signal between neutrophils and fibrin was identical in both conditions, with 100% of the fibrin generated colocalizing with neutrophils **(Supl. Figure 2D)**. Similarly, the expression of tissue factor on the surface of neutrophils recruited within the thrombus was equivalent between the two groups, supporting the statement that the initiation of coagulation induced by aircraft conditions requires mechanisms distinct from the extrinsic pathway (**Supl. Figure 3**). Taken together, these results indicate that neutrophils play a crucial role in activating coagulation in DVT, independent of changes in the atmospheric gas partial pressure.

**Figure 4.**
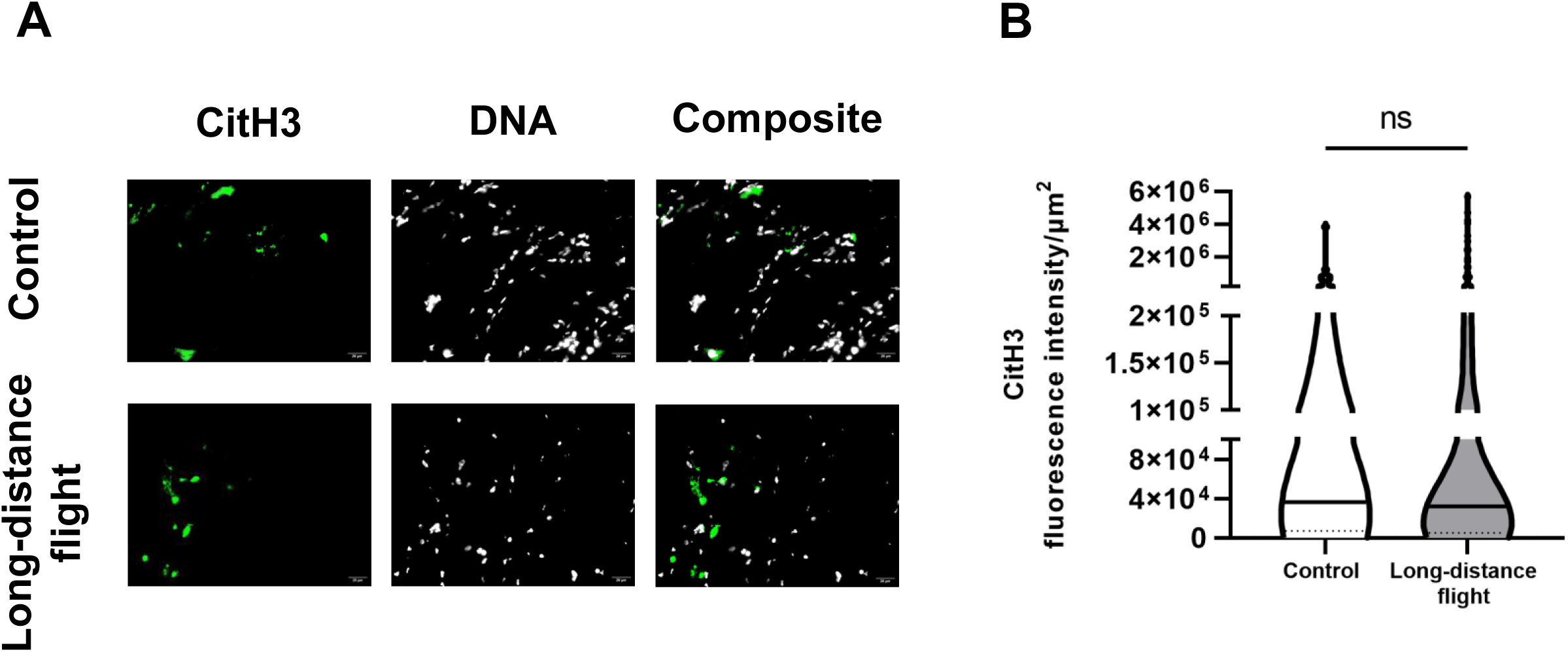
The pro-thrombotic effect during flight is independent of NETosis. **(A)** Representative images captured with x40 objective of CitH3 expression in thrombi formed in mice exposed to normobaric normoxia (control) and hypobaric hypoxia (long-distance flight) conditions. **(B)** Graphs depict the fluorescence intensity of CitH3 in both groups (n=5). Twenty-four hours post-IVC stenosis, thrombi were collected in both groups and fixed in OCT to perform cryostat sections (5 µm). NETs were detected with anti-CitH3 antibody labeled with Alexa Fluor 647 (1 µg/ml) and Hoechst 33342 for DNA labeling. (Mann-Whitney test).

**Figure 5.**
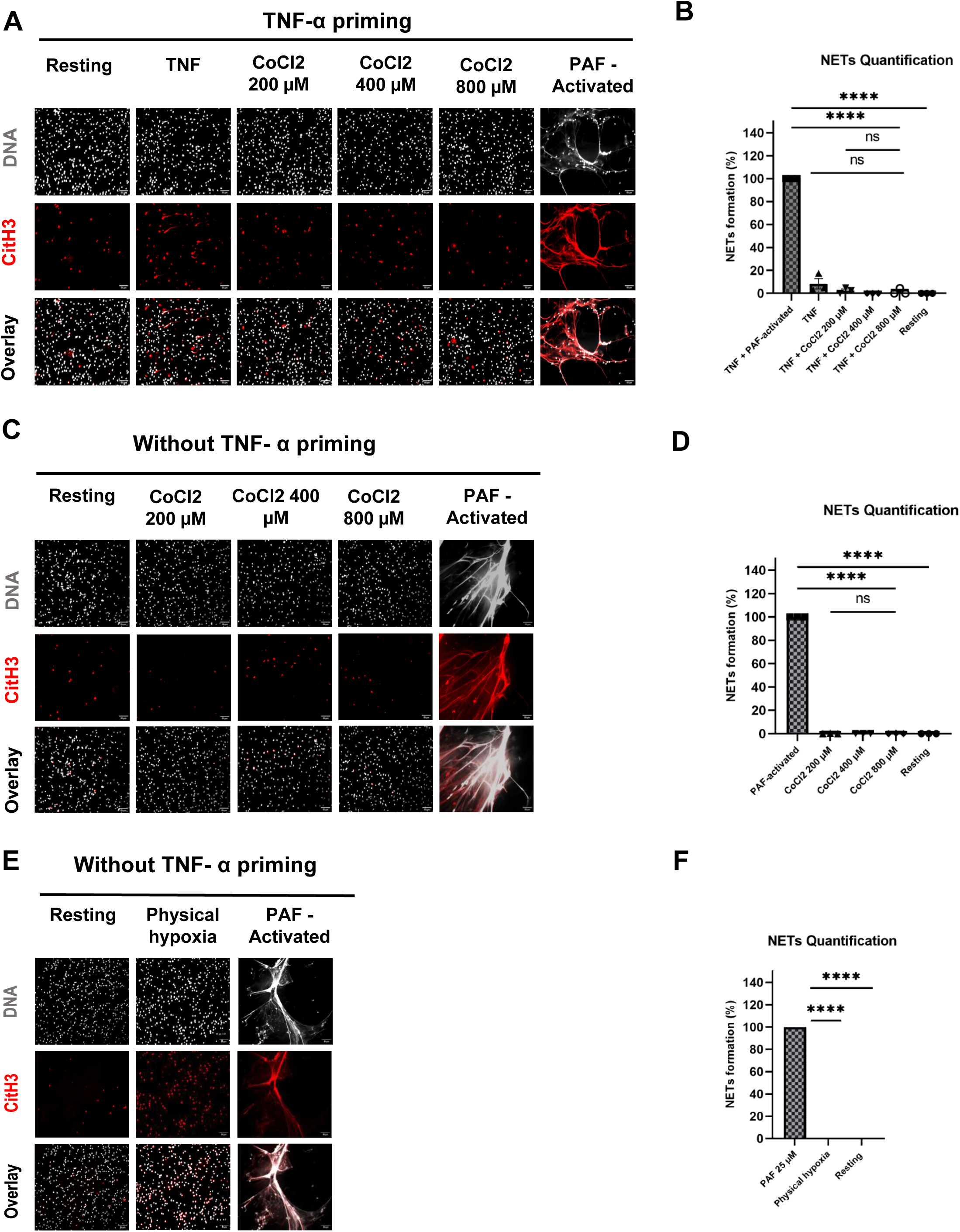
Hypoxia alone doesn’t induce NETs formation in vitro. **(A, C)** Representative images of CitH3 expression on murine neutrophils primed **(A)** or not **(C)** with TNF-alpha and treated with three concentrations of cobalt chloride during 3 hours. Neutrophils were immunostained with anti-CitH3 antibody (in red) and Hoechst 33342 for DNA labeling (in gray). Superposition of both labeling (DNA/CitH3+) was represented in pink. **(B, D)** Graphs depict the percentage of NETosis from murine neutrophils in each condition with **(B)** or without **(D)** TNF-alpha priming (mean+/- SEM). PAF-activated neutrophils condition was used as positive control corresponding to 100% of NETs formation (ANOVA test, **** P≤0,0001). **(E)** Representative images of CitH3 expression on murine neutrophils exposed to 4% of oxygen in hypoxia chamber without TNF-alpha priming. Neutrophils were immunostained with anti-CitH3 antibody (1,4 µg/ml) (in red) and Hoechst 33342 for DNA labeling (in gray). Superposition of both labeling (DNA/CitH3+) was represented in pink. **(F)** Graphs depict the percentage of NETosis from murine neutrophils in each condition without TNF-alpha priming (mean+/- SEM). PAF-activated neutrophils condition was used as positive control corresponding to 100% of NETs formation (ANOVA test, **** P≤0,0001).

### ADP-dependent platelets activation is important in the development of DVT during long-distance flight

We compared the colocalization between the platelet membrane receptor GPIb and P-selectin in the thrombus of control and exposed mice. Compared to control conditions, hypobaric hypoxia induced increased P-selectin expression at the surface of platelets recruited in the thrombus **(Figure 6A)**. Although the Pearson’s coefficient was identical between the two conditions, the area of colocalization of platelets and P-selectin was higher in mice exposed to long-distance flight conditions compared to control group **(Figure 6B and 6C)**. Furthermore, the proportion of P-selectin colocalized with platelets was higher after exposure to long-distance flight conditions compared to control group **(Figures 6D-F)**. Altogether, these results indicate that long-distance flight conditions induce an overactivation of platelets. ADP has been previously described to be generated after exposure to hypobaric hypoxia conditions (9). To confirm that ADP may be generated in circulation in our hypobaric hypoxia conditions, the concentration of adenosine, the end product of the degradation of purines, was determined. Hypobaric hypoxia led to an increase in circulating levels of adenosine up to 1,01±0,537 µM, compared with the control group with rates 7 times lower (0,138±0,038 µM) **(Figure 7A)**. Interestingly, the elevation of circulating adenosine concentration was also observed in the absence of IVC ligation **(Figure 7B)** and was independent of the surgery in the flight conditions, unlike the control group **(Figures 7C and 7D)**. The results indicate that the increased levels of adenosine were not induced by the inflammation caused by surgery but rather by the environmental conditions of the aircraft. To determine a potential involvement of ADP-dependent platelet activation in the pro-thrombotic effect of a long-distance flight, we next compared thrombus formation in *P_2_RY_12_ ^-/-^* mice and clopidogrel-treated mice. Interestingly, we no longer observed any morphological differences between thrombi developed in the two environmental conditions when platelets activation to ADP was inhibited, in contrast to the previous observations in wild type mice (**Figure 8A**). The weight of thrombi formed after DVT was identical between the two groups of mice, indicating that thrombi formed in clopidogrel-treated and *P_2_RY_12_ ^-/-^* mice were no longer affected by changes in the atmospheric gas partial pressure. (**Figure 8B**). Additionally, in contrast to wild-type mice exposed to hypobaric hypoxia conditions, where all of them experienced thrombosis, only 87,5% of *P_2_RY_12_ ^-/-^* mice and 83% of mice treated with clopidogrel developed thrombosis (**Figure 8C and 8D**). We conclude that the absence or the blocking of P_2_RY_12_ protected mice from the pro-thrombotic profile induced by mimicked long-distance flight conditions. When thrombus weight was compared between wild type and *P_2_RY_12_ ^-/-^* mice or clopidogrel-treated mice during flight, we observed that the weight of the thrombus was significantly decreased by up to 70% and 58,8 % respectively when platelet activation to ADP was inhibited **(Figure 8E)**. However, thrombus weight was not affected by the lack of P_2_RY_12_ receptor or treatment with clopidogrel in controls mice, suggesting that, unlike aircraft conditions, DVT development in a standard environment was independent of platelet activation to ADP **(Figure 8F)**.

**Figure 6.**
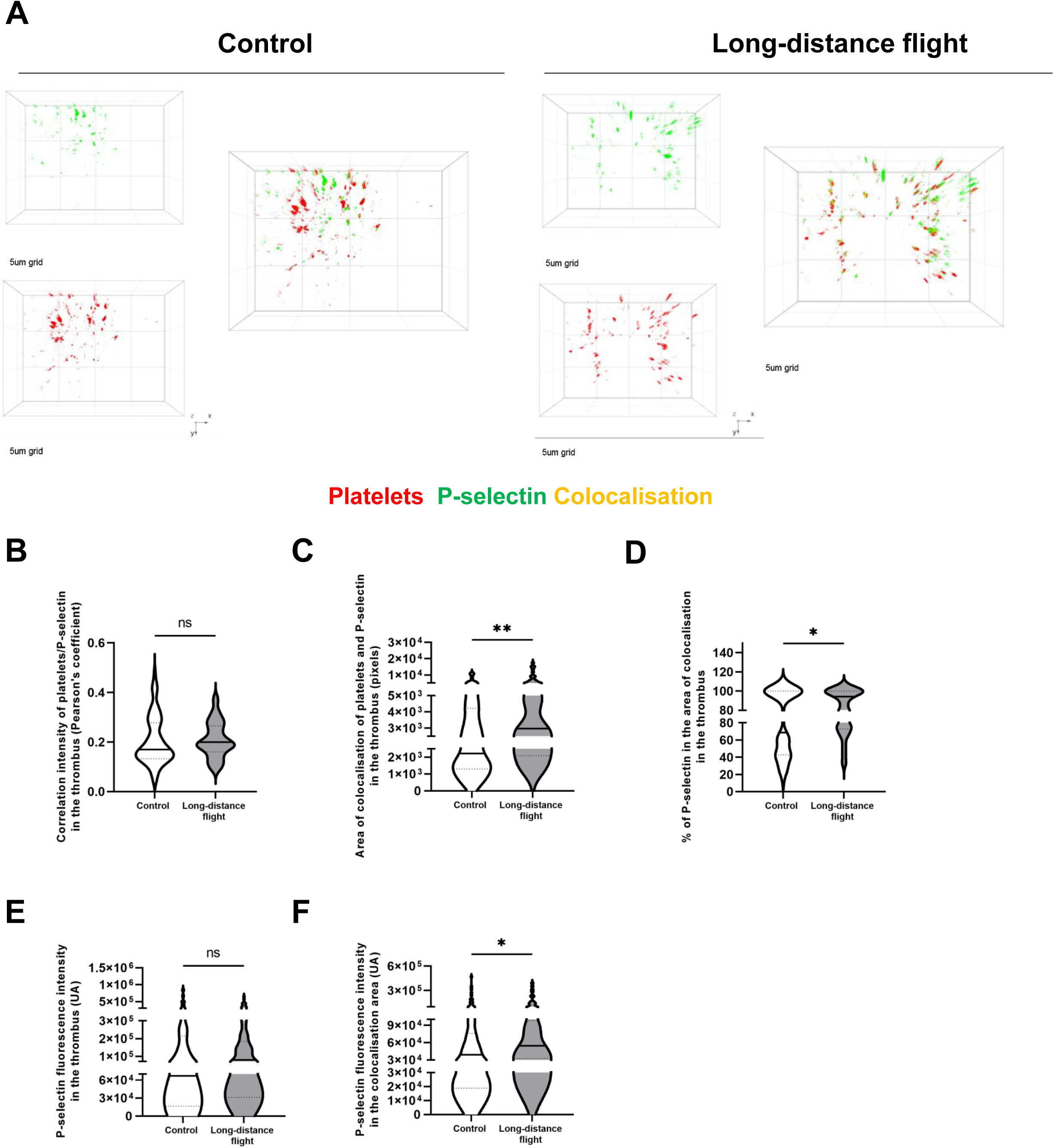
P-selectin expression on platelets is increased after exposure to long-distance flight conditions. **(A)** 3D Representative images captured with x60 water-immersion objective in confocal microscopy of platelets and P-selectin in mice exposed to normobaric normoxia (control) and hypobaric hypoxia (long-distance flight) conditions. **(B-D)** Graphs represent the Pearson’s correlation between the two signals **(B)**, the area of colocalization **(C)** and the percentage of P-selectin colocalized with platelets **(D)** under control or long-distance flight conditions. **(E-F)** Fluorescence intensity of total P-selectin **(E)** and of P-selectin in the area of colocalization **(F)** in thrombi of mice exposed to control or long-distance flight conditions (n=4). Twenty-four hours post-IVC stenosis, thrombi were collected in both groups and fixed in OCT to perform cryostat sections (5 µm). Platelets were detected with GPIb-beta antibody conjugated with Alexa Fluor 488 (1 µg/ml) and P-selectin with CD62P antibody conjugated with Alexa Fluor 647 (1 µg/ml). Lines show the median. (Mann-Whitney test, * P≤0,05, ** P≤0,005).

**Figure 7.**
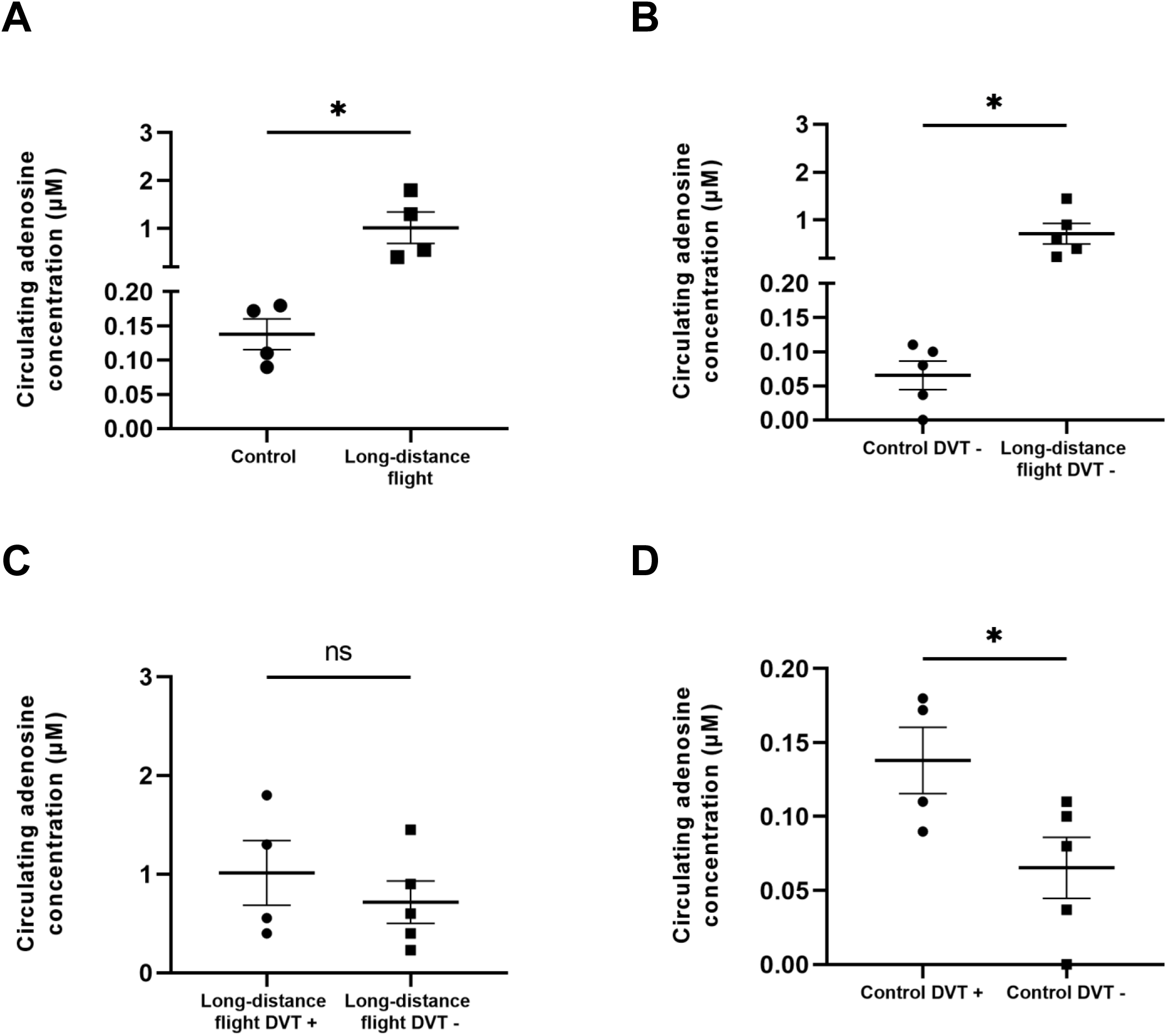
Long-distance flight conditions increase the circulating levels of adenosine. **(A-B)** Graphs depict the circulating concentration of adenosine in mice with **(A)** or without **(B)** the induction of inferior vena cava (IVC) stenosis under normobaric normoxia (control) and hypobaric hypoxia (long-distance flight) conditions. Comparison of adenosine levels with or without surgery under standard aircraft **(C)** or standard conditions **(D)**. (Student Test, *P≤0,05).

**Figure 8.**
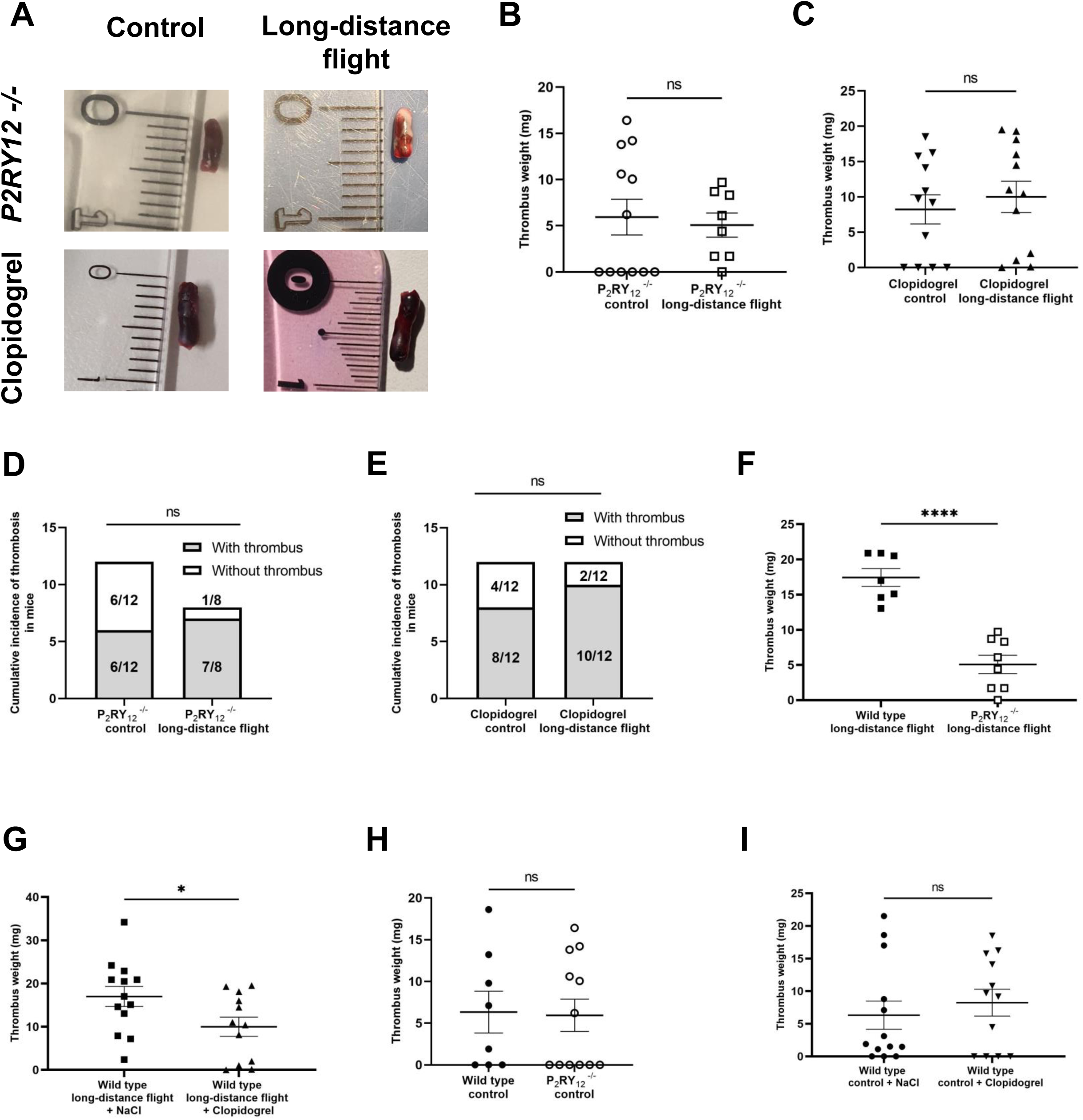
Long-distance flight conditions no longer affect thrombus weight when platelets activation to ADP is altered. **(A)** Representative pictures of thrombi collected in *P_2_RY_12_^-/-^* and clopidogrel-treated mice exposed to control or long-distance flight conditions. **(B-C)** Graph represents weight of thrombi in *P_2_RY_12_^-/-^* mice **(B)** and clopidogrel-treated mice **(C)** exposed to control or long-distance flight conditions. **(D-E)** Graphs illustrate the aggregated occurrence of thrombosis in *P_2_RY_12_^-/-^* mice **(D)** and clopidogrel-treated mice **(E)** 24 hours following the induction of inferior vena cava (IVC) stenosis under normobaric normoxia (control) and hypobaric hypoxia (long-distance flight) conditions. **(F-I)** Comparison of thrombus weight under aicraft **(F-G)** or control **(H-I)** conditions between wild type vs *P_2_RY^-/-^* mice and wild type vs clopidogrel-treated mice. (Student Test, * P≤0,5, ****P≤0,00005).

Taken together, these results delineate that prolonged air-travel exacerbates thrombus formation in DVT, primarily by altering the thrombus composition. This alteration notably manifests as an augmented recruitment of leukocytes and neutrophils. Intriguingly, the mechanisms contributing to this enhanced thrombus formation appear to be independent of NETs but dependent of ADP-activated platelets, opening a potential strategy for the prevention of venous thrombosis associated with long-distance flights.

## Discussion

Our results demonstrate that long-distance flight conditions disrupt hemostasis by inducing a pro-coagulant/pro-thrombotic profile that promote the development of DVT. Changes in the atmospheric gas partial pressure, generated in our experimental model, increase the recruitment of coagulation partners to the thrombus, as well as the secretion of ADP that activates platelets. We therefore observe that inhibition of platelets activation to ADP prevents the pro-thrombotic response generated by aircraft conditions, presenting platelets as key partners in the development of DVT during air travel.

Various clinical studies have reported an association between air travel and the development of DVT. The reduced oxygen and pressure conditions in the cabin have been described as the main cause for the development of VTE, and the occurrence of thrombosis increases with the duration of the flight as well as with the pre-existence of cardiovascular risk factors (10–13). In our study, mice exposed to 6 hours of hypobaric hypoxia develop larger and denser thrombi than the control group. Our experimental model of a long-distance flight reproduces and supports the pro-thrombotic effect observed clinically, allowingus to investigate the mechanism involved in the development of DVT following air travel.

The flow restriction model of DVT in the IVC is commonly used and is described as reproducing the kinetics and the clinical features of DVT in humans by reducing venous flow (14). Using this model, in combination with intravital imaging, von Brühl et al. characterized the thrombus by a successive layer of red blood cells and fibrin, called “red clot”, associated with a large number of leukocytes and activated platelets (3). Mechanical stress and local hypoxia generated by an impaired blood flow were described as disturbing the vasoprotective function of endothelial cells by promoting their activation and the release of the Weibel-Palade bodies (WPBs). Degranulation of the activated endothelium enhances the expression of P-selectin, leading to the rolling of neutrophils before their firm adhesion through the complexes ICAM-1/LFA-1 and VCAM-1/VLA-4 (15,16). The exocytosis of the WPBs and therefore the expression of pro-adhesive molecules by endothelial cells was described as being enhanced during hypoxia (17). In our study, we show that mice exposed to aircraft conditions formed thrombi richer in leucocytes and neutrophils, supporting the involvement of neutrophils in DVT and the deregulation of the vasoprotective properties of endothelium in these environmental conditions. The reduction of the circulating concentration of neutrophils observed in the flight group may therefore be explained by a greater local consumption.

Once recruited in the clot, neutrophils participate in thrombus formation by expressing pro-coagulant proteins from granules on their surface, such as NE and cathepsin G which inhibit TFPI and therefore trigger the activation of coagulation (18). Although several studies suggest that hypoxia stimulates the activation and degranulation of neutrophils, we did not observe an up-regulation of activation markers, either on circulating neutrophils or on those recruited to the thrombus. Nevertheless, studies indicating that reduced oxygen levels upregulate neutrophils degranulation were performed in vitro, under normobaric conditions, and at levels of hypoxia that do not reflect the conditions found during air travel (19).

Neutrophils are widely described in DVT due to their capacity to form NETs that activate the coagulation cascade by expressing serine proteases or promoting the binding of coagulation factors to their adhesive surface (20). However, the demonstration of the involvement of NETs in thrombus formation is mainly based on the use of DNAse that also degrades ATP and ADP, two potent platelet and neutrophil agonists. We previously demonstrated that the inhibition of thrombus formation by DNAse-1 could be independent of NETosis, suggesting that DNAse infusion was not sufficient to confirm the role of NETs in thrombosis and that other approaches should be considered (21). The impact of hypoxia on NETosis is controversial and reveals distinct results depending on the NETosis stimulus and the mode of hypoxia applied (physical or chemical). For instance, treatment of neutrophils with an iron chelator desferrioxamine (DFO) leads to NETs formation while incubation of human neutrophils for 5 hours under physical hypoxia totally abolished the PMA-induced NETosis (22,23). In our study, exposure of mouse neutrophils to either physical or chemical hypoxia did not induce NETs formation in vitro. Moreover, even in the pressure-reduced conditions in addition to hypoxia, no NETs are detected within thrombi developed in vivo. This would therefore suggest that the development of DVT during flight is independent of NETosis. Nevertheless, the whole fibrin generated in thrombi colocalized with neutrophils, under the two environmental conditions. Neutrophils are therefore the main cells involved in activating of coagulation in DVT, independent of NETs formation and regardless of environmental conditions. Although this colocalization did not increase under hypobaric hypoxia, the density of neutrophils was higher in thrombi developed during flight, resulting therefore in a greater supply of fibrin. The pro-coagulant function of neutrophils is well described in the DVT model as well as in arterial thrombosis, mainly through the expression of TF(15,24). Shut *et al.* revealed that the activation of coagulation during air travel was independent of the extrinsic pathway, which is in agreement with our observations (8). Although the hypothesis of fibrin generation by neutrophils via the intrinsic pathway is the most likely, further experiments are needed to identify the pathway involved in the activation of the coagulation cascade during DVT, both under standard or aircraft conditions (15,24).

The recruitment of platelets to the vessel wall is mediated by the interaction between the GPIb-α subunit and the vWF stored in WPBs and released by activated endothelial cells (25). Once adhered to the endothelium, platelets are activated by various agonists such as thrombin (PAR-1/PAR-4), ADP (P_2_RY_1_/P_2_RY_12_) and TxA2 (RTxA2), leading to calcium influx that enhances platelets degranulation and activates the integrin αIIbβ3 required for aggregation with fibrinogen. The first layer of activated platelets at the site of thrombus formation then provides the recruitment of other circulating platelets to promote the propagation of DVT. Although the recruitment of circulating elements to the vessel wall was increased during hypoxia, we observed no effect of aircraft conditions on the density of platelets within the thrombus, as well as on their circulating concentration. These results are consistent with the literature that suggests an increase in platelet count in response to altitude only over several days, correlated with a transcription of thrombopoietin which probably doesn’t occur during a flight of a few hours (26). In contrast, one clinical study showed a 20% reduction in the number of circulating platelets on the first day after a stay at 4 559m of altitude, in association with a retention in the pulmonary capillaries (27). However, although the platelet density in thrombi was identical between the two environmental conditions, platelets in thrombi developed in mice exposed to hypobaric hypoxia express a greater amount of P-selectin and are therefore more activated. Indeed, several studies have reported that high-altitude conditions induce platelet hyperreactivity through an increase of platelet aggregation in response to ADP, fibrinogen or collagen, as well as an elevation in the expression of αIIbβ3 (9,28,29). In addition, circulating levels of ATP and ADP, which are platelets agonists via their purinergic receptors, are increased when oxygen levels are reduced, as confirmed by measurement of circulating adenosine concentration in mice exposed to aircraft conditions. The rise in circulating ADP levels during flight then stimulates platelets via P_2_RY_1_/P_2_RY_12_ receptors, promoting therefore the activation of coagulation and the development of DVT.

RBCs are a potent source of ATP (1-5 mM) and its secretion is stimulated by shear stress, mechanical deformation and hypoxia. These cells are the efficient Hg-containing carriers of oxygen and therefore adapt to circulating oxygen depletion to maintain tissue supply. For this purpose, erythrocytes adjust their carbohydrate metabolism by activating the glycolysis pathway, inducing a high production of ATP that stimulates NO generation to increase blood flow and therefore maintain O_2_ supply in hypoxia (30). ATP could be released from RBCs via transporters, mainly via the Panx-1 channel, or by hemolysis which is the favored hypothesis because it requires less energy than active transport (31). Moreover, ATP modulates the submembrane spectrin network and could therefore induce vesiculation of RBCs, leading to their lysis. We observed fewer RBCs within thrombi developed in mice exposed to aircraft conditions, supporting the hemolysis hypothesis. Conversely, some studies demonstrated an increase in RBCs production under reduced oxygen levels due to erythropoietin synthesis mediated by HIF1-alpha (32,33). However, transcription of erythropoietin, and the subsequent erythropoiesis, requires more than 6 hours and is therefore not achieved on a long-distance flight. The ATP secreted by RBCs is then degraded to ADP by ectonucleotidase (CD39/CD73), whose expression is also increased in hypoxia, allowing therefore the stimulation of platelets. Altogether, these results indicate that ADP-dependent platelets activation is important for the development of DVT during flight and open up new therapeutic approaches by using antiplatelets instead of or in combination with anticoagulants to prevent these coagulation disorders.

## Methods

### Mice

Wild Type C57BL/6J (5 to 12 weeks) mice were obtained from Janvier-Labs. Homozygous *P_2_RY_12_ ^-/-^* mice were originally from MRC Harwell and were bred at CEFOS for all the experiments.

### Antibodies and reagents

A rat anti-mouse LY6G antibody labeled with PE (clone 1A8, BD Biosciences), a rat anti-mouse GPIb-beta Dylight 649 (clone X649, Emfret analytics) and a rat anti-mouse Ter-119 labeled with PE (clone Ter-19, BD Biosciences) were used for both labelling of circulating cells and to study the thrombi composition. A mouse anti-human CD11b activated labeled with FITC (clone CBRM1/5, eBioscience) and elastin (EnzChek Elastase Assay Kit, ThermoFisher) were used in association with a rat anti-mouse LY6G antibody for the evaluation of neutrophils activation in cytometry. To identify the circulating cellular composition of blood, a brilliant-violet 421 nm rat anti-mouse CD45 (clone 30-F11, Biolegend), a rat anti-mouse Ly6C labeled with PC7 (clone AL-21, BD Biosciences) and a rat anti-mouse CD16/CD32 (clone 2.4G2, BD Biosciences) were used A rat anti-mouse CD45 coupled with APC (clone 30-F11, Ozyme), a rabbit anti-mouse CitH3 (clone ab5103, Abcam), a mouse anti-human tissue factor (clone TF9-10H10, Abcam), a rabbit anti-mouse HIF1alpha (Proteintech) and a labeled mouse anti-fibrin antibody were used as previously described (34). Alexa Fluor 647-conjugated anti-rabbit IgG1 (clone MOPC-21, Biolegend) was used as a secondary antibody when necessary. The rat anti-mouse GPIb-beta Dylight 488 (clone X488, Emfret analytics) and the rat anti-mouse CD62P Alexa Fluor 647 (clone RB40.34, BD Biosciences) were used to performed confocal microscopy experiments.

### Mouse model of flow restriction in the IVC

Stenosis in the IVC was created using the DVT model as described by Holly Payne and Alexander Brill (14) and adapted to our specific condition (hypobaric and hypoxia). 24 hours before the stenosis procedure, the animals were acclimatized to the water intake with a gel. Briefly, mice were previously intraperitoneally injected with buprenorphine (0,1 mg/kg) and anesthetized with isoflurane during surgery. A median laparotomy was performed to move a part of digestif tract outside the abdominal cavity and expose the IVC. During all the surgery the intestinal tract were superfused with 37°C of saline buffer. All visible side branches below the bifurcation of the left renal vein were completely ligated with 5:0 propylene suture to control venous stasis and improve reproducibility. After separation from aorta below the bifurcation of the left renal vein, the IVC was ligated with 7:0 polypropylene suture with a spacer placed over the vessel to create a 90% partial occlusion. The spacer was then gently removed to avoid endothelial injury. Once the ligature performed, peritoneum and skin were closed separately with 4:0 propylene sutures. After waking up, the animals were placed in a cage with access to water and buprenorphine in a gel. Mice were euthanized after 24 hours and thrombi developed in the IVC were collected for further analysis. Sham experiments consisted to move a part of the bowel out of the abdominal cavity without performed any ligations.

### Mice exposure to simulated long-distance flight conditions

Wild type C57BL/6J and *P_2_RY ^-/-^* mice were exposed for six hours to simulated long-distance flight conditions to an altitude of 2000 meters corresponding to a pO2 of 0.8 bars (i.e 115 mmHg) in a home designed hypobaric chamber that remained at room temperature. The animals had access to gelled water including buprenorphine and food during the simulated flight, as under control conditions.

### Mice treatment

Wild type C57BL/6J were orally treated with clopidogrel (8 mg/kg, dilution in NaCl 0,9%) two days before the DVT surgery and the following day to maintain the reduction of platelet activation to ADP until thrombus recovery.

### Flow cytometry

Analyses of CD45, Ly6G, Ly6C, Ter 119 and GPIb-beta expression on the membrane of circulating cells were performed by flow cytometry (Gallios, Beckman Coulter).

### Circulating blood cell phenotyping

Wild type C57BL/6J mice were anesthetized with isoflurane and previously injected intraperitoneally with buprenorphine (0,1 mg/kg). Blood was collected from intracardiac sampling on citrated buffer 24 hours after IVC ligation or sham experiments. For erythrocytes and platelets detection, blood samples were incubated for 20 minutes at room temperatures with CD16/CD32 to prevent non-specific signal before antibodies incubation (a rat anti-mouse Ter-119 labeled with PE and a rat anti-mouse GPIb-beta labeled with Alexa fluor 647) for 30 min at room temperature. To detect leucocytes, a lysis of red blood cells was previously performed for 10 minutes at room temperature. Then, leukocytes were incubated with a brilliant-violet 421 nm rat anti-mouse CD45, a rat anti-mouse LY6G antibody labeled with PE and a rat anti-mouse Ly6C labeled with PC7for 30 minutes at 4°C to ovoid their activation. Counting beads (Beckman coulter) were added in all samples to assess cell concentration.

### Prothrombin time assay

Whole blood was collected from the tail of mice 24 hours after IVC ligation or sham experiments. The quantitative prothrombin time (PT) was evaluated with the Xprecia Stride Coagulation System. Briefly, blood sample was applied to a test strip and drawn by capillary action into the reaction chamber of the strip were a mix of reagents activates the coagulation cascade. The test stops when the blood is clotted and the results is displayed in seconds or International Normalized Ratio (INR).

### Evaluation of circulating adenosine concentrations

Whole blood (20 was collected from the tail of mice 24 hours after IVC ligation or sham experiments and dried on blotting paper (Whatman 903 protein saver cards™). Then, the blot spots were stored at room temperature, until analysis. Blood Samples extraction and analyze have been previously reported (35). Briefly, six mm of blood spot were cut out followed by extraction (mix of internal standard and methanol) for 90 min at 45 °C. Then aliquots were evaporated to dryness at 60 °C under nitrogen; 150 μL of 0.1% formic acid in water were added and quickly vortexed before transferring into an HPLC auto sampler vial. Samples were analyzed using a Shimadzu UFLC XR system (Shimadzu, Marne la Vallee, France). The LC system was interfaced with an ABSciex 4500 triple quadrupole mass spectrometer (Shimatzu, Les Ulis, France) operating with an electrospray ionization source (ESI) using nitrogen (purity: 99.99%). 10 µL of the extracted sample were injected onto a 2.1 × 100 mm, 3 μm column (Waters, Guyancourt, France). The mobile phase was 3% methanol and 97% acidified water (0.1% formic acid) with a flow of 0.7 mL/min for 3.5 min. The gradient of methanol was increased to 30% for 3 min.

### Immunohistochemistry from thrombi sections

OCT-thrombi sections (5 µm) were obtained with the Leica CM1900 Cryostat (Leica) for immunolabelling of cellular and protein composition. Briefly, after permeabilization with cold methanol for 5 minutes, thrombi sections were blocked with 3% of BSA for 30 minutes in a humidity chamber. Thrombi were then incubated with appropriate primary antibodies for 1h30 minutes. After successive washes in PBS^+/+^, secondary antibodies were added for 45 minutes when necessary. Nuclei were labeled with Hoechst for 10 minutes and mounting medium was added on the thrombi section after extensive washes in PBS ^+/+^ and secured with a lamella (ProLong Gold Antifade, Thermofisher Scientific). Thrombi sections were visualized with the EVOS cell imaging system (Thermo Fisher Scientific) minimum 24 hours after the assembling of the slide and lamella. Immunofluorescence data were obtained using an Olympus 40x objective. For each section, we captured as many images as possible, without returning to the same location, to obtain an overview of the thrombus. Image analysis was performed using ImageJ and the integrated density of fluorescence was assessed for each capture. The fluorescence intensity of markers was analyzed by determining median integrated density values of all thrombi sections.

### Confocal microscopy

Immunohistochemistry on OCT-thrombi sections were performed as previously described. Confocal imaging of thrombi sections was obtained via a spinning disk (Intelligent Imaging Innovations, Denver, CO) and immunofluorescence data were obtained using an Olympus AX microscope with a 60x water-immersion objective. Image analysis was performed using Slidebook 6 (Intelligent Imaging Innovations) to evaluate the Pearson’s correlation coefficient and the area of colocalization of the two markers.

### Purification of mouse polymorphonuclear neutrophils

The extraction and purification of murine neutrophils was performed as previously described by Darbousset et al (21).

### In vitro NETs experiments

Purified neutrophils were primed or not with 2 ng/ml of TNF-alpha during 30 minutes and then incubated for 1 hour on 24-well plates coated with L-polylysine. NETs formation was induced for 3 hours at 37°C and 5% CO2 by stimulation with 10 µM of PAF (Calbiochem) on primed neutrophils and 25 µM on non-primed ones. Resting neutrophils remained in Hanks ^+/+^ without agonists as a negative control of NETosis. For neutrophils exposed to oxygen deprivation, they were incubated for 3 hours at 37°C and exposed in a hypoxic chamber to a normobaric hypoxic atmosphere obtained after N2 flushing (1% O2, 5% CO2 and 94% N2) or in presence of cobalt chloride (CoCl2 at 200, 400 and 800 µM), a chemical way to stabilize the hypoxia-inducible factor 1 alpha subunit in order to mimic hypoxia signaling. At the end of this process, supernatant in each well was removed and neutrophils were fixed with 4% paraformaldehyde for 10 minutes at room temperature. To perform immunofluorescence, neutrophils were incubated for 1h30 minutes with an anti-citrullinated histone 3 (4°C). Then, after extensive washings, neutrophils were incubated with a secondary antibody conjugated to Alexa Fluor 647 during 45 minutes in the dark (4°C). Nuclei were labeled with Hoechst for 10 minutes in the dark at room temperature. Neutrophils and NETs were visualized under a fluorescence microscope (EVOS cell imaging system, Thermo Fisher Scientific).

### Statistical analysis

For the in vivo experiments, the significance was determined using unpaired two-tailed Student’s t test or unpaired Mann-Whitney U test when the samples did not follow a normal distribution. The ex vivo experiments were analyzed using unpaired Mann-Whitney U test and for the in vitro experiments the significance was determined using an Ordinary one-way ANOVA. Differences were considered significant at P≤0,05.

### Study approval

All animal care and experimental procedures were performed as recommended by the European Community guidelines and approved by the Marseille ethical comity #14 (APAFIS#23640-2020011614211724 and APAFIS#36998-202208291446686).

## Data availability

Data available on request from the authors.

## Author contributions

Contribution: J.T., E.C, L.C., L.B. and N.A. contributed to the design and performance of the research, analysis of data, and writing of the manuscript; and J.T., R.G., C.D. and L.P-D. contributed to the experimental design, analysis of data, and writing of the manuscript.

## Acknowledgments

We acknowledge Stephane Robert (Center for CardioVascular and Nutrition Research, Aix Marseille University) for his contribution to the optimization of phenotyping in flow cytometry.

---

**Figure Supl. 1.**
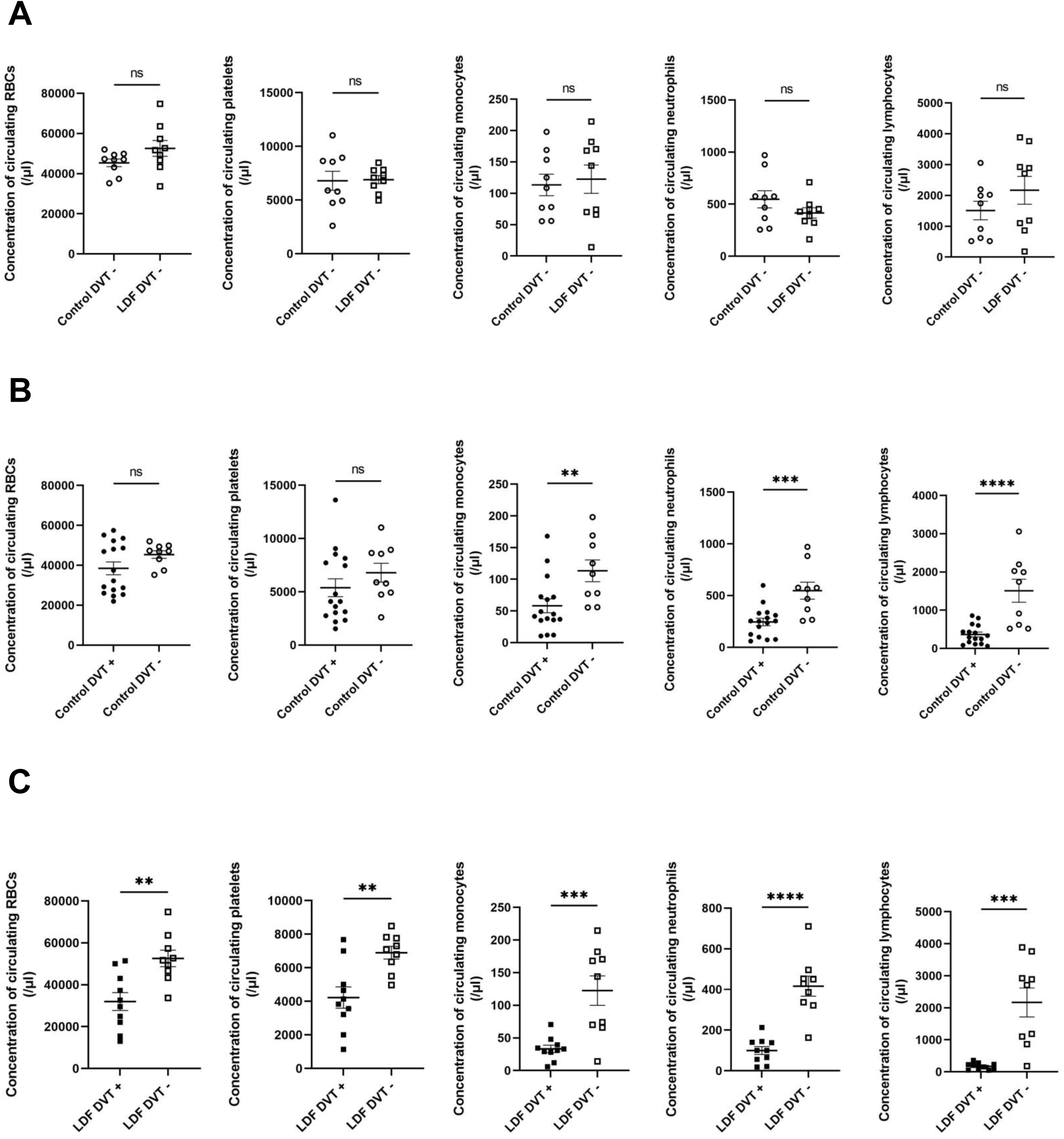
Local inflammation generated by the surgery decreased the count of leucocytes in circulation. **(A)** Graphs represent the circulating concentration of erythrocytes, platelets, monocytes, neutrophils and lymphocytes in the two environmental conditions without IVC ligation. **(B-C)** Graphs depict the circulating concentration of erythrocytes, platelets, monocytes, neutrophils and lymphocytes in mice exposed to standard **(B)** or aircraft **(C)** conditions with and without IVC ligation. Twenty-four hours post-IVC stenosis, blood samples were collected, and the circulating concentration of cells was assessed using flow cytometry. Erythrocytes were labeled with anti-Ter119 antibody (0,5 µg/ml), platelets with anti-X488 antibody (0,1 µg/ml), neutrophils with anti-Ly6G antibody (0,2 µg/ml), monocytes with anti-Ly6C antibody (0,2 µg/ml) and lymphocytes were detected as the CD45^+^, Ly6G^-^ and LY6C^-^ population. (Student Test, **P≤0,005, ***).

**Figure Supl. 3.**
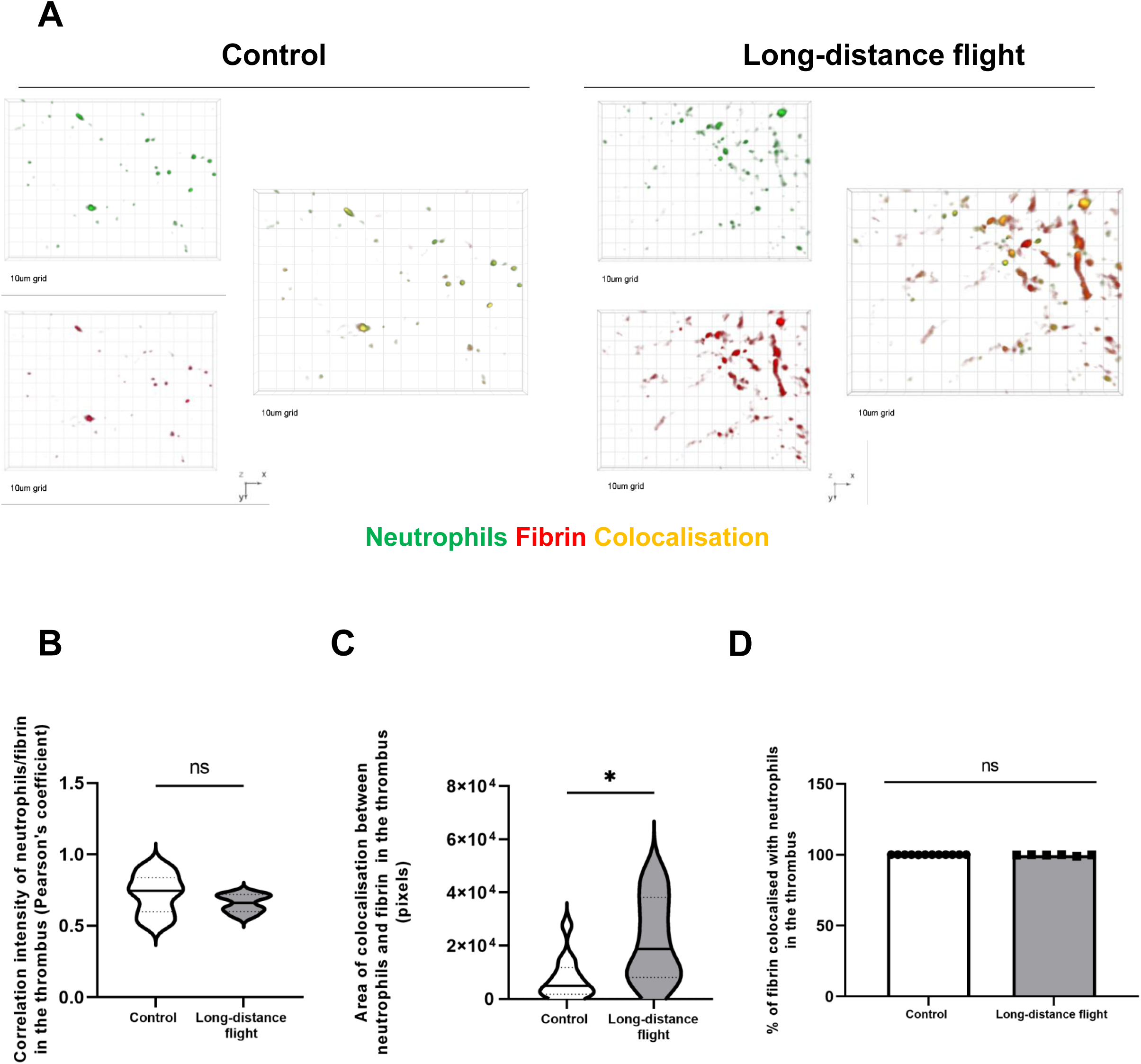
The extrinsic pathway is not involved in thrombus formation during a long-distance flight. **(A)** 3D representative images captured in confocal microscopy with x60 water-immersion objective of neutrophils and tissue factor in thrombi thrombi of mice exposed to control or aircraft conditions. **(B-D)** Graph represent the Pearson’s correlation between the two signals **(B),** the area of colocalization (C) and the percentage of tissue factor **(D)** colocalized with neutrophils in both conditions (n=4). Twenty-four hours post-IVC stenosis, thrombi were collected in both groups and fixed in OCT to perform cryostat sections (5 µm). Neutrophils were detected with anti-Ly6G antibody conjugated with PE (1 µg/ml) and fibrin with anti-fibrin antibody (1 µg/ml). Lines show the median. (Mann-Whitney test).

**Figure Supl 3.**
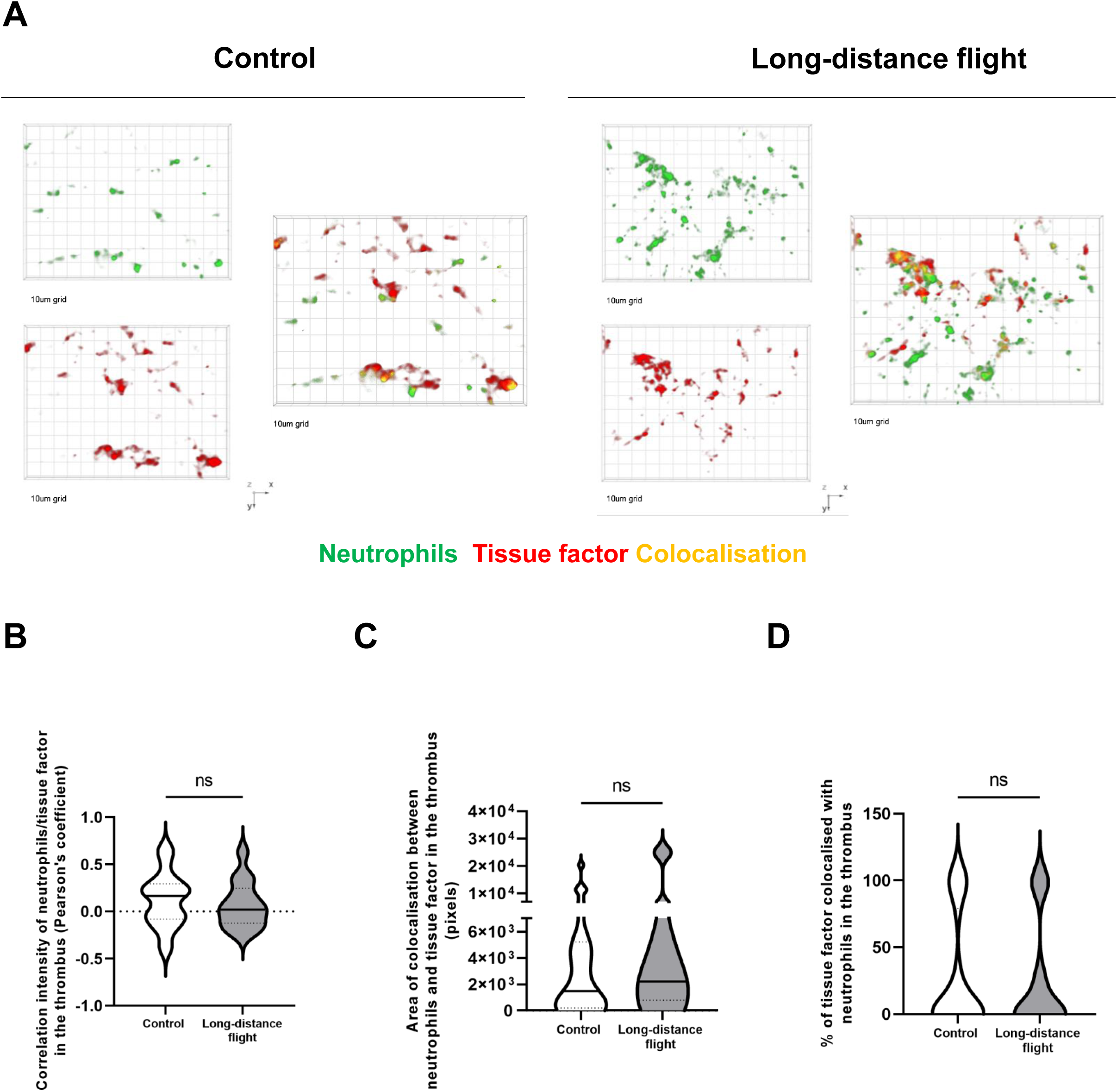
The extrinsic pathway is not involved in thrombus formation during a long-distance flight. **(A)** 3D representative images captured in confocal microscopy with x60 water-immersion objective of neutrophils and tissue factor in thrombi thrombi of mice exposed to control or aircraft conditions. **(B-D)** Graph represent the Pearson’s correlation between the two signals **(B),** the area of colocalization (C) and the percentage of tissue factor **(D)** colocalized with neutrophils in both conditions (n=4). Twenty-four hours post-IVC stenosis, thrombi were collected in both groups and fixed in OCT to perform cryostat sections (5 µm). Neutrophils were detected with anti-Ly6G antibody conjugated with PE (1 µg/ml) and tissue factor with anti-tissue factor antibody conjugated with Alexa Fluor 647 (1 µg/ml). (Mann-Whitney test).

## References

1. Khan F, et al. Venous thromboembolism. The Lancet. 3 juill 2021;398(10294):64-77.

2. Kumar DR, et al. Virchow’s Contribution to the Understanding of Thrombosis and Cellular Biology. Clin Med Res. déc 2010;8(3-4):168-72.

3. Mackman N. New insights into the mechanisms of venous thrombosis. J Clin Invest. juill 2012;122(7):2331-6.

4. Symington IS, Stack BH. Pulmonary thromboembolism after travel. Br J Dis Chest. avr 1977;71(2):138-40.

5. Schreijer AJM, et al. Activation of coagulation system during air travel: a crossover study. Lancet Lond Engl. 11 mars 2006;367(9513):832-8.

6. Krasiński Z, et al. COVID-19, long flights, and deep vein thrombosis: What we know so far. Cardiol J. 2021;28(6):941-53.

7. Schobersberger W, et al. Changes of biochemical markers and functional tests for clot formation during long-haul flights. Thromb Res. oct 2002;108(1):19-24.

8. Schut AM, et al. Coagulation activation during air travel is not initiated via the extrinsic pathway. Br J Haematol. juin 2015;169(6):903-5.

9. Shang C, et al. The human platelet transcriptome and proteome is altered and pro-thrombotic functional responses are increased during prolonged hypoxia exposure at high altitude. Platelets. 2020;31(1):33-42.

10. Watson HG. Travel and thrombosis. Blood Rev. sept 2005;19(5):235-41.

11. Watson HG, Baglin TP. Guidelines on travel-related venous thrombosis. Br J Haematol. janv 2011;152(1):31-4.

12. Chee YL, Watson HG. Air travel and thrombosis. Br J Haematol. sept 2005;130(5):671-80.

13. Kuipers S, et al. The risk of venous thrombosis after air travel: contribution of clinical risk factors. Br J Haematol. mai 2014;165(3):412-3.

14. Payne H, Brill A. Stenosis of the Inferior Vena Cava: A Murine Model of Deep Vein Thrombosis. J Vis Exp. 22 déc 2017;(130):56697.

15. von Brühl ML, et al. Monocytes, neutrophils, and platelets cooperate to initiate and propagate venous thrombosis in mice in vivo. J Exp Med. 9 avr 2012;209(4):819-35.

16. Yago T, et al. Cooperative PSGL-1 and CXCR2 signaling in neutrophils promotes deep vein thrombosis in mice. Blood. 27 sept 2018;132(13):1426-37.

17. Pinsky DJ, et al. Hypoxia-induced exocytosis of endothelial cell Weibel-Palade bodies. A mechanism for rapid neutrophil recruitment after cardiac preservation. J Clin Invest. 15 janv 1996;97(2):493-500.

18. Massberg S, et al. Reciprocal coupling of coagulation and innate immunity via neutrophil serine proteases. Nat Med. août 2010;16(8):887-96.

19. Hoenderdos K, et al. Hypoxia upregulates neutrophil degranulation and potential for tissue injury. Thorax. nov 2016;71(11):1030-8.

20. Fuchs TA, et al. NET impact on deep vein thrombosis. Arterioscler Thromb Vasc Biol. août 2012;32(8):1777-83.

21. Carminita E, et al. DNAse-dependent, NET-independent pathway of thrombus formation in vivo. Proc Natl Acad Sci. 13 juill 2021;118(28):e2100561118.

22. Völlger L, et al. Iron-chelating agent desferrioxamine stimulates formation of neutrophil extracellular traps (NETs) in human blood-derived neutrophils. Biosci Rep. juill 2016;36(3):e00333.

23. Branitzki-Heinemann K, et al. Formation of Neutrophil Extracellular Traps under Low Oxygen Level [published online November 25, 2016]. *Front Immunol.* 10.3389/fimmu.2016.00518

24. Darbousset R, T et al. Tissue factor-positive neutrophils bind to injured endothelial wall and initiate thrombus formation. Blood. 6 sept 2012;120(10):2133-43.

25. Brill A, et al. von Willebrand factor-mediated platelet adhesion is critical for deep vein thrombosis in mouse models. Blood. 27 janv 2011;117(4):1400-7.

26. Hartmann S, et al. Effect of altitude on thrombopoietin and the platelet count in healthy volunteers. Thromb Haemost. janv 2005;93(1):115-7.

27. Lehmann T, et al. Platelet count and function at high altitude and in high-altitude pulmonary edema. J Appl Physiol (1985). févr 2006;100(2):690-4.

28. Rocke AS, et al. Thromboelastometry and Platelet Function during Acclimatization to High Altitude. Thromb Haemost. janv 2018;118(1):63-71.

29. Tyagi T, et al. Altered expression of platelet proteins and calpain activity mediate hypoxia-induced prothrombotic phenotype. Blood. 20 févr 2014;123(8):1250-60.

30. Grygorczyk R, Orlov SN. Effects of Hypoxia on Erythrocyte Membrane Properties-Implications for Intravascular Hemolysis and Purinergic Control of Blood Flow. Front Physiol. 2017;8:1110.

31. Sikora J, et al. Hemolysis is a primary ATP-release mechanism in human erythrocytes. Blood. 25 sept 2014;124(13):2150-7.

32. Kuhrt D, Wojchowski DM. Emerging EPO and EPO receptor regulators and signal transducers. Blood. 4 juin 2015;125(23):3536-41.

33. Haase VH. Regulation of erythropoiesis by hypoxia-inducible factors. Blood Rev. janv 2013;27(1):41-53.

34. Palacios-Acedo AL, et al. P2RY12-Inhibitors Reduce Cancer-Associated Thrombosis and Tumor Growth in Pancreatic Cancers. Front Oncol. 2021;11:704945.

35. Groppelli A, et al. Adenosine Concentration in Patients With Neurally Mediated Syncope. Front Cardiovasc Med. 2022;9:900023.

